# Reducing neuronal nitric oxide synthase (nNOS) expression in the ventromedial hypothalamus (VMH) increases body weight and blood glucose levels while decreasing physical activity in female mice

**DOI:** 10.1101/2024.05.15.594324

**Authors:** Pamela R. Hirschberg, Dashiel M. Siegel, Suraj Teegala, Munir Rahbe, Pallabi Sarkar, Vanessa H. Routh

## Abstract

Glucose-inhibited (GI) neurons of the ventromedial hypothalamus (VMH) depend on neuronal nitric oxide synthase (nNOS) and AMP-activated protein kinase (AMPK) for activation in low glucose. The Lopez laboratory has shown that the effects of estrogen on brown fat thermogenesis and white fat browning require inhibition of VMH AMPK. This effect of estrogen was mediated by downstream lateral hypothalamus (LH) orexin neurons^1,2^. We previously showed that estrogen inhibits activation of GI neurons in low glucose by inhibiting AMPK^3^. Thus, we hypothesized that VMH AMPK- and nNOS-dependent GI neurons project to and inhibit orexin neurons. Estrogen inhibition of AMPK in GI neurons would then disinhibit orexin neurons and stimulate brown fat thermogenesis and white fat browning, leading to decreased body weight. To test this hypothesis, we reduced VMH nNOS expression using nNOS shRNA in female mice and measured body weight, adiposity, body temperature, white and brown fat uncoupling protein (UCP1; an index of thermogenesis and browning), locomotor activity, and blood glucose levels. Surprisingly, we saw no effect of reduced VMH nNOS expression on body temperature or UCP1. Instead, body weight and adiposity increased by 30% over 2 weeks post injection of nNOS shRNA. This was associated with increased blood glucose levels and decreased locomotor activity. We also found that VMH nNOS-GI neurons project to the LH. However, stimulation of VMH-LH projections increased excitatory glutamate input onto orexin neurons. Thus, our data do not support our original hypothesis. Excitation of orexin neurons has previously been shown to increase physical activity, leading to decreased blood glucose and body weight^4^. We now hypothesize that VMH nNOS-GI neurons play a role in this latter function of orexin neurons.

## INTRODUCTION

We showed previously that ∼60% of neurons in the ventrolateral ventromedial hypothalamus (vlVMH) are glucose-inhibited (GI) neurons that depend on AMP-activated protein kinase (AMPK) to sense glucose^3,5^. GI neurons isolated from the entire VMH require neuronal nitric oxide synthase (nNOS) and AMPK to sense glucose and over 75% of VMH neurons which produce NO are GI neurons^5,6^. Thus, there is almost complete overlap between VMH nNOS neurons and AMPK-dependent GI neurons. Lopez and colleagues found that VMH AMPK inhibition is required for brown adipose tissue (BAT) thermogenesis and white adipose tissue (WAT) browning in response to estrogen^1,7^. This effect of estrogen was mediated by lateral hypothalamus (LH) orexin neurons which are downstream of estrogen sensitive VMH AMPK inhibition^2^. We found that estrogen blunts activation of vlVMH GI neurons in low glucose by inhibiting AMPK^3^. Based on these observations, we originally hypothesized that 1) vlVMH AMPK-dependent GI neurons are also dependent on nNOS and 2) inhibition of these neurons mediated the stimulatory effect of estrogen on LH orexin neurons, BAT thermogenesis and WAT browning.

To test our hypothesis, we first confirmed that vlVMH neurons were NOS dependent and that vlVMH-GI and -nNOS neurons project to the LH. Next, we reduced VMH nNOS expression in female mice using shRNA injection and determined whether BAT thermogenesis and WAT browning were increased due to reduced inhibition by VMH nNOS-GI neurons. Lastly, we determined whether VMH to LH projections modulated synaptic input onto orexin neurons. Interestingly, our data did not support our original hypothesis. VMH nNOS shRNA had no effect on BAT thermogenesis, WAT browning, body temperature or serum free fatty acid (FFA) and epinephrine levels. However, VMH nNOS shRNA increased body weight by 30% in 14 days. This was associated with increased blood glucose and decreased locomotion. Finally, the majority of VMH to LH projections increased glutamatergic currents onto orexin neurons. Kotz and colleagues have shown that LH orexin neurons lower blood glucose by increasing spontaneous physical activity^4^. Together, these data suggest the revised hypothesis that VMH nNOS-GI neurons activate LH orexin neurons, leading to increased locomotion, decreased blood glucose and a reduction in body weight.

## METHODS

### Animals

Male and female 6 to 10-week-old C57BL/6J mice, mice expressing green fluorescent protein (GFP) in orexin neurons (orexin-GFP mice; bred in house^8^), mice expressing cre-recombinase in nNOS neurons (nNOS-cre mice; Jackson Laboratories B6.129-*Nos1^tm1(cre)Mgmj^*/J # 017526) or mice expressing cre-dependent Zsgreen (B6.Cg-*Gt(ROSA)26Sor^tm6(CAG-ZsGreen1)Hze^*/J # 007906) were housed on a 12:12 light-dark cycle with food and water provided ad libitum. Naïve mice were group housed and mice following stereotaxic surgery were housed singly. All procedures were approved by the Rutgers Newark Institutional Animal Care and Use Committee (IACUC).

### Adeno-associated virus (AAV) and Retrobeads™

The following were obtained from commercial vendors: shRNA nNOS (AAV1-GFP-U6-m-NOS1-shRNA, Vector Biolabs, Malvern PA), scrambled control (AAV1-GFP-U6-scrambled-shRNA, Vector Biolabs, Malvern PA), Channelrhodopsin (ChR2) expressing virus (pAAV9-hSyn-hChR2(H134R)-mCherry) (AddGene, 26976-AAV9, Watertown, MA) and Retrobeads™ (Red RetroBeads, Lumaflor, Durham, NC)

### Stereotactic surgery

Mice were anesthetized with ketamine (80-100mg/kg)/ Xylazine (100mg/mL) i.p. and analgesic (Buprenorphine SR, 1mg/kg) was given subcutaneously. A borosilicate pipette tip was pulled to a length of at least 7mm and back filled with mineral oil. Retrobeads™ or AAV were front filled and injected using a *Nanoject* (Drummond Scientific) attached to the stereotaxic frame (Kopf Instruments, Tujunga CA). Topical anesthetic (Bupivacaine) was injected under the skin of the head prior to surgery. Bregma was set as point zero using a Digital Display Readout Console (940-C, Kopf instruments). Lateral hypothalamus coordinates: A/P: -1.30 M/L: +/-.95 and D/V. - 5.05 and the VMH coordinates A/P: -1.33 M/L: +/-55 and D/V: -5.65. Retrobeads™ were injected unilaterally and AAV were injected bilaterally.

### Electrophysiology

Male and female C57BL/6J or orexin-GFP were anesthetized with sodium pentobarbital (60–80 mg/kg, i.p.) and transcardially perfused with ice-cold oxygenated (95%O2/5%CO2) N-methyl- d -glucamine (NMDG) perfusion solution (composition in mM: 110 NMDG, 2.5 KCl, 1.25 NaH _2_ PO _4_, 30 NaHCO _3_, 20 N-2-hydroxyethylpiperazine-N′-2-ethanesulfonic acid (HEPES), 10 glucose, 2 thiourea, 0.5 CaCl _2_, 10 MgSO _4_ ·7H _2_ O, 5 Na-ascorbate, 3 Na-pyruvate (pH 7.3– 7.4, osmolarity adjusted to 310–315 mOsm). Brains were rapidly removed, placed in ice-cold (slushy) oxygenated NMDG perfusion solution and 300 μm coronal slices containing VMH neurons were made on a vibratome (7000 smz2, Vibroslice, Camden Instruments, Camden, UK) as previously described^9^. The brain slices were then transferred to a pre-warmed (34 °C) initial recovery chamber filled with 150 ml of NMDG perfusion solution. After transferring the slices, Na ^+^ was reintroduced following the Na ^+^ -spike method^10^. Slices were then transferred to HEPES-artificial cerebrospinal fluid (aCSF) holding solution (composition in mM: 92 NaCl, 2.5 KCl, 1.25 NaH _2_ PO _4_, 30 NaHCO _3_, 20 HEPES, 2.5 glucose, 2 thiourea, 5 Na-ascorbate, 3 Na-pyruvate, 2 CaCl _2_ ·2H _2_ O, and 2 MgSO _4_ ·7H _2_ O (pH 7.3–7.4, 310–315 mOsm) and allowed to recover for 1 h at room temperature prior to whole-cell current or voltage clamp recording using the recording aCSF (composition in mM: 124 NaCl, 2.5 KCl, 1.25 NaH _2_ PO _4_, 24 NaHCO _3_, 2.5 glucose, 5 HEPES, 2 CaCl _2_ ·2H _2_ O, and 2 MgSO _4_ ·7H _2_ O (pH 7.3–7.4, 310–315 mOsm).

Data were collected using using pClamp v11.3 software and an AxoPatch 200B amplifier and digitized with a DigiData 1500B AD converter (Molecular Devices). Neurons were visualized using infrared differential interference contrast and/or fluorescence microscopy. aCSF was perfused at a rate of 3-5mL/minute. Borosilicate pipettes were pulled to 3-5mOhm resistance and back filled with pipette solution (current clamp: K^+^-Gluconate 128mM, EGTA 0.5mM, KCl 10mM, HEPES 10mM, MgCl2. 6H20 4mM, CaCl2 2H20 0.5mM, osmolarity 290-300 mOsm, pH 7.1-7.2; voltage clamp: KCl 125mM, KGluconate 10mM, MgCl2 2mM, EGTA 0.2mM, HEPES 10mM, Mg2 ATP 2mM, Na2 GTP 0.5mM; osmolarity 290-300 mOsm, pH 7.1-7.2). Voltage clamp recordings during optogenetic experiments were made at either 0mV or -60mV. Whole cell current clamp recordings are used to identify glucose sensing neurons. A 300msec -10 or -20pA hyperpolarizing current was injected every 3 seconds. GI neurons were identified by a reversible depolarization and increase in cellular resistance as glucose decreases from 2.5 to 0.1 mM. The voltage change to the injected current was calculated for 10 consecutive voltage responses at the height of the response to low glucose. For sGI, these ten responses were taken from the very end of the 10-minute recording. For adGI neurons the 10 responses at the height of the neuron response were averaged. Ohm’s law (V=IR) is used to calculate the resistance based on the value of injected current in Amps (I) and the average voltage response in Volts (V). The change in cell resistance was required to be at least 15% in response to low glucose to characterize the neuron as GI.

### In vitro optogenetics

Orexin or non-orexin neurons in th*e* LH were located by visualizing GFP or lack thereof, respectively and recorded in the voltage clamp configuration (-60 mV holding potential). ChR2 expressing neurons were excited using a 50msec flash of 470nm blue light. Holding neurons at -60mV with a high K+ internal solution (recipe) yielded reversal potentials of 128mV for Na+, - 120mV for K+ and 0mV for Cl-). Thus, the K+ current is outward, while Cl- and Na+ currents are inward. 3mM kynurenic acid (blocker of all glutamatergic channels) was used to block glutamate currents, and 100uM saclofen (GABA_B_ antagonist) and 200mM bicuculline (GABA_A_ antagonist) to block GABA currents. The neurons are exposed to inhibitors for at least five minutes before recordings begin. A response to the VMH neuron stimulation is determined by a response during the blue light duration in at least 75% of the total sweeps (10) in each recording.

### Western Blot

BAT, subcutaneous WAT, and VMH samples were immediately put in ice cold cell extraction buffer (recipe 6). After sonication and BAT/WAT samples were centrifuged at 10,000 RPG for 15 min at 4 degrees C. and VMH samples at 12,000 RPM for 20 min at 4 degrees. The supernatant was then collected and stored for protein quantification and gels. Total protein content of BAT, subcutaneous WAT, and VMH protein lysates were determined using a BCA protein assay (Thermo Fisher). Samples were probed with antibodies against UCP1 (Thermo Fisher, PA1-24894, 1:5000), or nNOS (Cell Signaling, C7D7 1:1000), and then Β actin (Sigma, A1978, 1:2000). Chemiluminescent imaging was done with Immobilon ® Classico Western HRP Substrate (Millipore Sigma WBLUC). Resultant bands were quantitated using Image J software (NIH, Bethesda, Maryland, USA). Protein was expressed in relation to β actin levels.

### Serum free fatty acid and epinephrine quantification

Mice were sacrificed under anesthesia by exsanguination through intra-cardiac blood draw with a 3mL syringe affixed with a 25 gauge needle. Whole blood was immediately dispensed into serum separator tubes (MiniCollect® TUBE 0.8 ml, Greiner 450472) and left to clot at room temperature for two hours. Blood was then centrifuged at 1,000g for 20 minutes at 4 degrees C. Serum was separated and frozen at -80 before use in either free fatty acid (AbCam cat no. ab65341) or epinephrine (Adrenaline ELISA Kit, AbCam, cat no. ab287788) quantification assays.

### IHC (brain)

*30µM sections:* Animals were perfused with 4% paraformaldehyde (PFA). Brains were kept for 24hr in 4% PFA, cryoprotected with 15 and 30% sucrose solution in 0.01M phosphate buffered saline (PBS) and stored in -80°C in OCT (tissue tek, Sakura). After sectioning 30uM sections, the tissue was blocked with 10% BSA in PBS for one hour. Post blocking, the slices were probed with primary antibody. Tissue was washed 3 times for 10 min in PBS containing Tween 20 before placing in secondary antibody. Slices were mounted on slides with mounting media containing nuclear counterstain (ProLong™ Gold Antifade Mountant with DNA Stain DAPI, Thermo Fisher P36931) and cover glass. *300µM section IHC*: Following recording, slices were fixed in 4% PFA for 48 hours, Slices were then incubated in primary antibody for 48hours at 4C and in secondary for 1 hour. All images were acquired using a fluorescent microscope equipped with confocal-like capabilities.

### Immunohistochemistry (IHC) (fat)

Both inguinal white fat and subcutaneous brown fat samples were fixed in 4% PFA for four days before transferring to 70% ethanol. Fat samples were paraffin embedded, stained with UCP1 antibody, and analyzed by the Rutgers Histology Core.

### Cell Counting

All cell counting was performed manually on ImageJ software using the cell counting feature.

### Quantifying physical activity and interscapular temperature

After stereotaxic injections, RFID/temperature probes (Unified Identification Devices (UID), Lake Villa, IL) are injected into the interscapular region. The RFID chips allow for temperature information when read with handheld reader (URH-1HP, UID), or when mouse cage is placed on the Mouse Matrix Home Cage Monitoring System (UID). Placement on Mouse Matrix allows for fast tracking of movement between 8 cage zones as well as temperature. Activity index is calculated based on “zone to zone movements”. To calculate the activity, the distance between zones is measured from one antenna center to another using the X axis as 3.625” and the Y axis as 3.16”. The total distance from one recorded scan to another is calculated via the use of the Pythagorean Theorem (a^2^ + b^2^ = c^2^).”

### Statistical analysis

All statistical analyses were performed using GraphPad Prism v8. Comparisons between groups were made using an independent t-test and within groups using a paired t-test. The values were expressed as the means ± SEM. The differences with p<0.05 were considered statistically significant.

## RESULTS

As we have shown previously, two subpopulations of vlVMH GI neurons were observed. “Sustained” (s) GI neurons remained depolarized in response to decreased glucose from 2.5 to 0.1 mM until glucose levels returned to 2.5 mM whereas “adapting” (adGI) neurons returned to baseline membrane potential while the neurons are still exposed to 0.1 mM glucose. Of 25 vlVMH GI neurons 15/25 were sGI and 10/25 were adGI (Table 1). These two subsets of GI neurons did not differ in average resistance (R) or membrane potential (MP) in 2.5 or 0.1mM glucose. Further, they are not different in percent change in MP (unpaired *t-test t(23)=1.19, mean ±SEM: sGI 8.26±1.21 vs. adGI 11.52±2.84, p=0.25*) or R (unpaired t-test: t(23)=1.48, *mean ±SEM: sGI 47.46± 5.35 vs. adGI 69.13± 16.17, p=0.15)* when glucose is lowered from 2.5 to 0.1mM. However, adGI neurons take significantly less time to respond to decreased glucose (unpaired t-test, *t(23)=6.35*, n=15, *mean ± SEM*: sGI 602.9*±23.82 vs. adGI 310.8± 43.81*, p<0.0001).

**Table 1.** Comparison of the electrophysiological properties of ventrolateral ventromedial hypothalamus (vlVMH) sustained (s) glucose-inhibited (GI) and adjusting (ad) GI neurons. A. Average resistance (R) and membrane potential (MP) in 2.5 mM and 0.1 mM glucose glucose, percent change in R and MP as glucose decreased from 2.5 to 0.1 mM and the time to the peak response for each GI subtype. B. Graphs representing the averages presented in A.

| Average values | R 2.5mM (MΩ) | R 0.1mM (MΩ) | MP 2.5mM (mV) | MP 0.1mM (mV) | % change MP | % change R | Time to peak response (s) |
| --- | --- | --- | --- | --- | --- | --- | --- |
| sGI (15) | 504.0 ± 50.1 | 733.7 ± 71.0 | -65.4 ± 2.5 | -60.2 ± 2.6 | 8.3 ± 1.2 | 47.5 ± 5.4 | 602.9 ± 23.8*** |
| adGI (10) | 485.5 ± 70.0 | 790.6 ± 103.3 | -60.0 ± 6.9 | -53.3 ± 2.7 | 11.5 ± 2.8 | 69.1 ± 16.2 | 310.8 ± 43.8*** |

### NOS is necessary for vlVMH GI neuron activation in low glucose

Although we found previously that neurons isolated from the entire VMH depend on nNOS to activate in low glucose^5^, it was important to verify that vlVMH GI neurons specifically reflect the rest of the VMH GI neuron population. We found that GI neurons of the vlVMH were also dependent on NOS to activate in low glucose. Application of two NOS inhibitors (L-NAME 100uM, L-NMMA 200uM, Tocris) to both sGI and adGI neurons (figure 1 A, B) led to a significant decrease in the percent change in R *(paired t-test, n=8, t(7)=3.451, mean ± SEM: control 69.80± 20.22 vs. NOS inhibitor 7.69± 3.19,* p=0.011)* and MP (*paired t-test t(10)=3.93, n=11*, *mean ± SEM: control 10.37±2.58 vs. NOS inhibitor 2.59±0.82, **p=0.003)* in response to lowering glucose from 2.5mM to 0.1mM.

**Figure 1.**
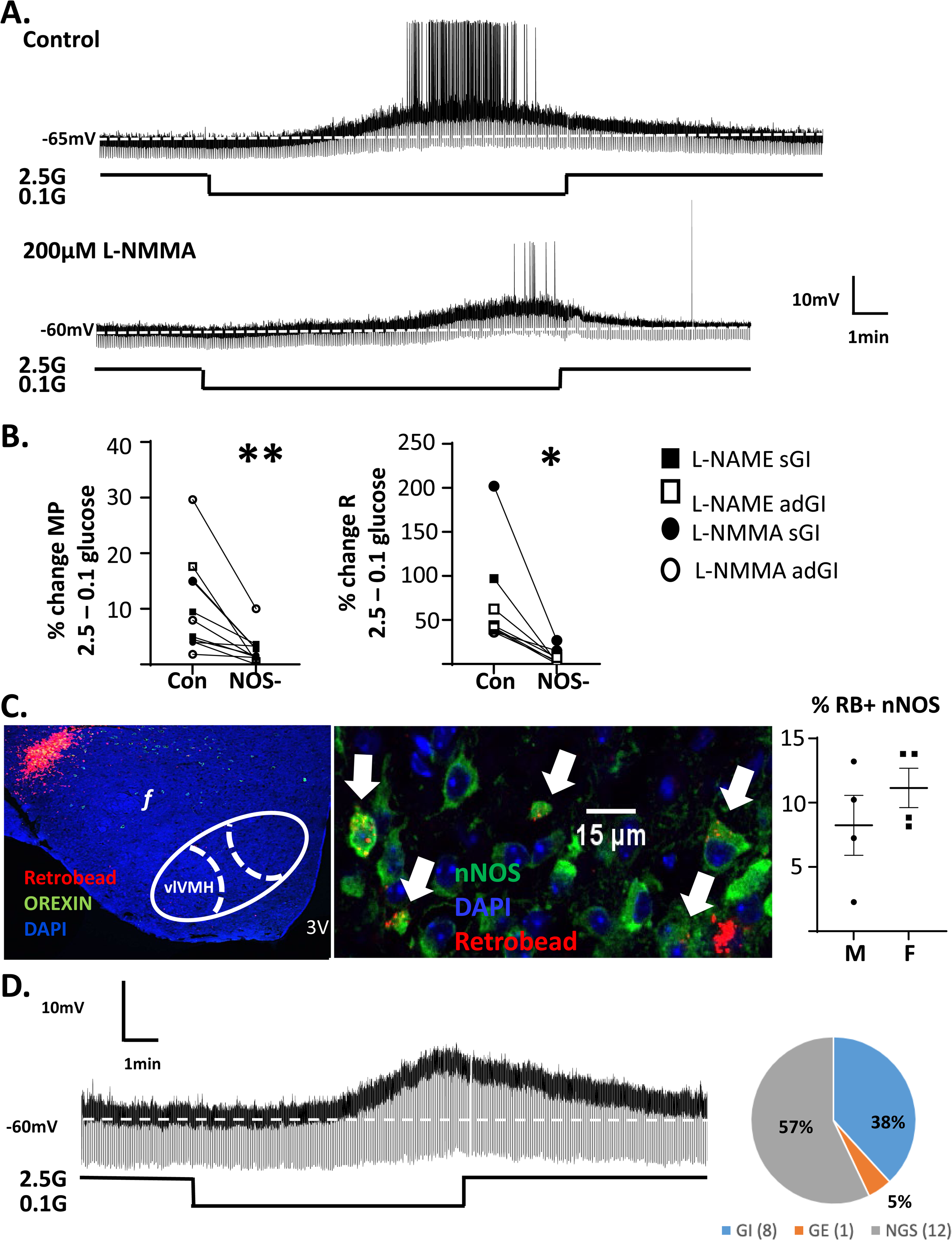
NOS mediates the response of vlVMH GI neurons to low glucose and vlVMH nNOS-GI neurons project to the LH. A. Representative current clamps traces of a vlVMH adGI neuron f in a brain slice. Lowering glucose from 2.5 to 0.1 mM in the presence of 200 μM L-NMMA (bottom trace) blunted the control response (top trace) of this neuron to decreased glucose. B. Graph of the percent change of MP and R for sGI and adGI neurons in the presence and absence of the NOS inhibitors, L-NAME or L-NMMA, as glucose decreased from 2.5 to 0.1 mM glucose. sGI neurons: black squares (L-NAME) or circles (L-NMMA); adGI: white squares or circles. C. Left panel: site of retrobead injection (red) in the orexin (green) field, blue – DAPI nuclear stain. Right panel: Retrobeads (red) taken up by nNOS neurons (green) in the vlVMH, blue – DAPI nuclear stain. Graph represents the percent of retrobead positive nNOS neurons in male (M) and female (F) mice. D. Representative trace of a retrobead positive sGI neuron exhibiting the typical sustained depolarization and increased R in low glucose. Pie chart represents the % GI, GI and non-glucose sensing retrobead positive neurons.

Next, we used fluorescently labeled Retrobeads^TM^ to determine whether vlVMH-nNOS and -GI neurons projected to the LH. Retrobeads were injected into the LH of male and female mice and retrobead positive vlVMH neurons were evaluated using IHC and electrophysiology. Retrobeads were observed in ∼10% of nNOS neurons of both sexes (Figure 1C). Moreover, approximately 40% (8 of 21) retrobead positive neurons were GI neurons, while only 1 retrobead positive neuron was a glucose-excited (GE) neuron (Figure 1D).

### VMH nNOS shRNA increases body weight and blood glucose levels

We first attempted to validate the commercially available nNOS-cre transgenic line from Jackson Laboratories (Strain #071526) to use opto- and/or chemogenetics to manipulate nNOS neurons. However, cre expression was not specific to nNOS neurons (Figure 2A-C). Therefore, we used shRNA to knock down nNOS expression in the VMH (Figure 3 A, B). We then examined the effects of VMH nNOS knockdown on measures of energy homeostasis.

**Figure 2.**
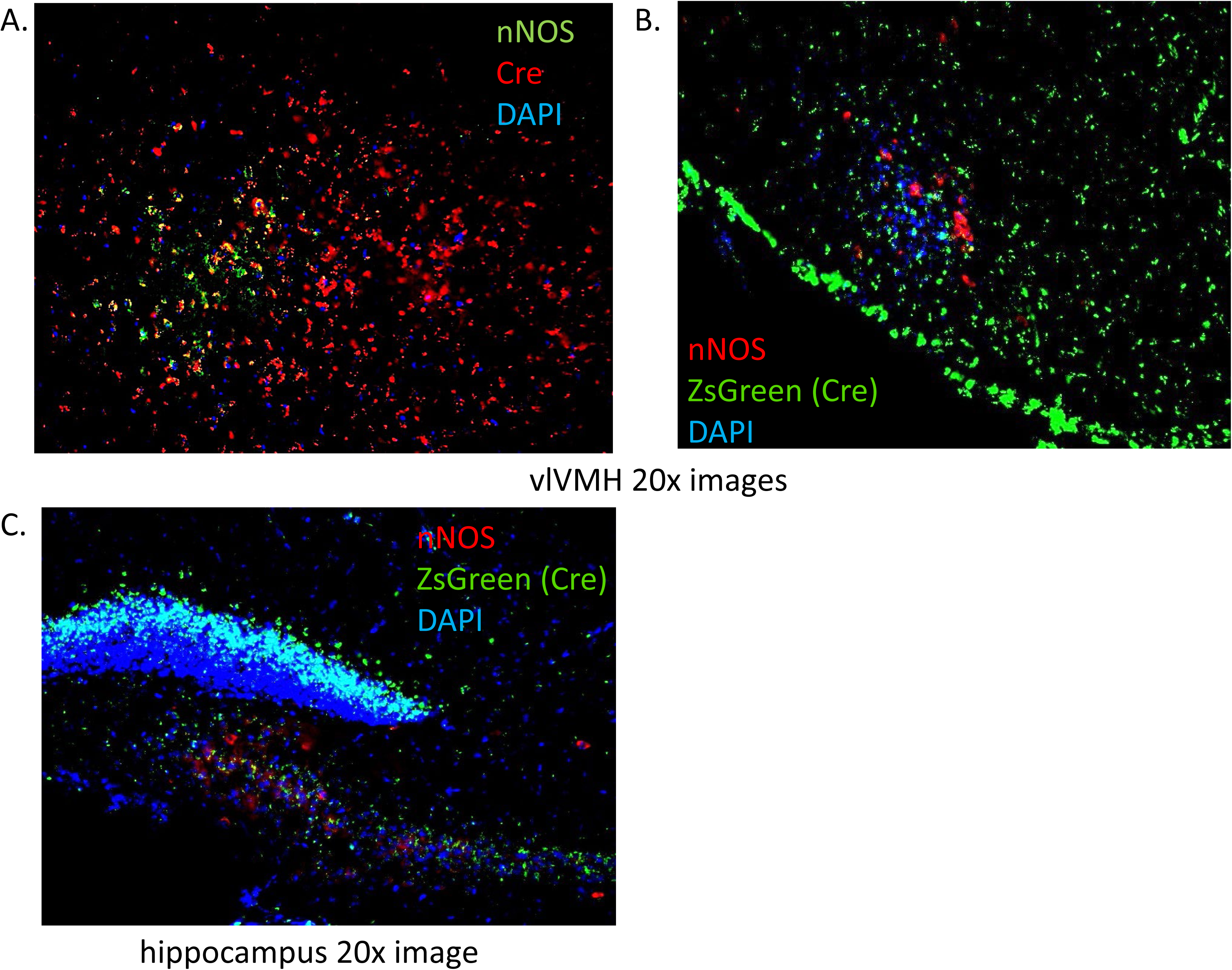
cre-recombinase expression was not specific to nNOS expressing neurons. A. cre-recombinase (red) was expressed in both vl VMH nNOS (green) and non-NOS expressing neurons; blue: DAPI neuronal marker. B, C. nNOS-cre mice were bred to mice expressing the cre-dependent fluorescent marker Zs-green. Zs-green was widely expressed in the vl VMH (B) and hippocampus (C) and was not specific to nNOS expressing neurons (red).

**Figure 3.**
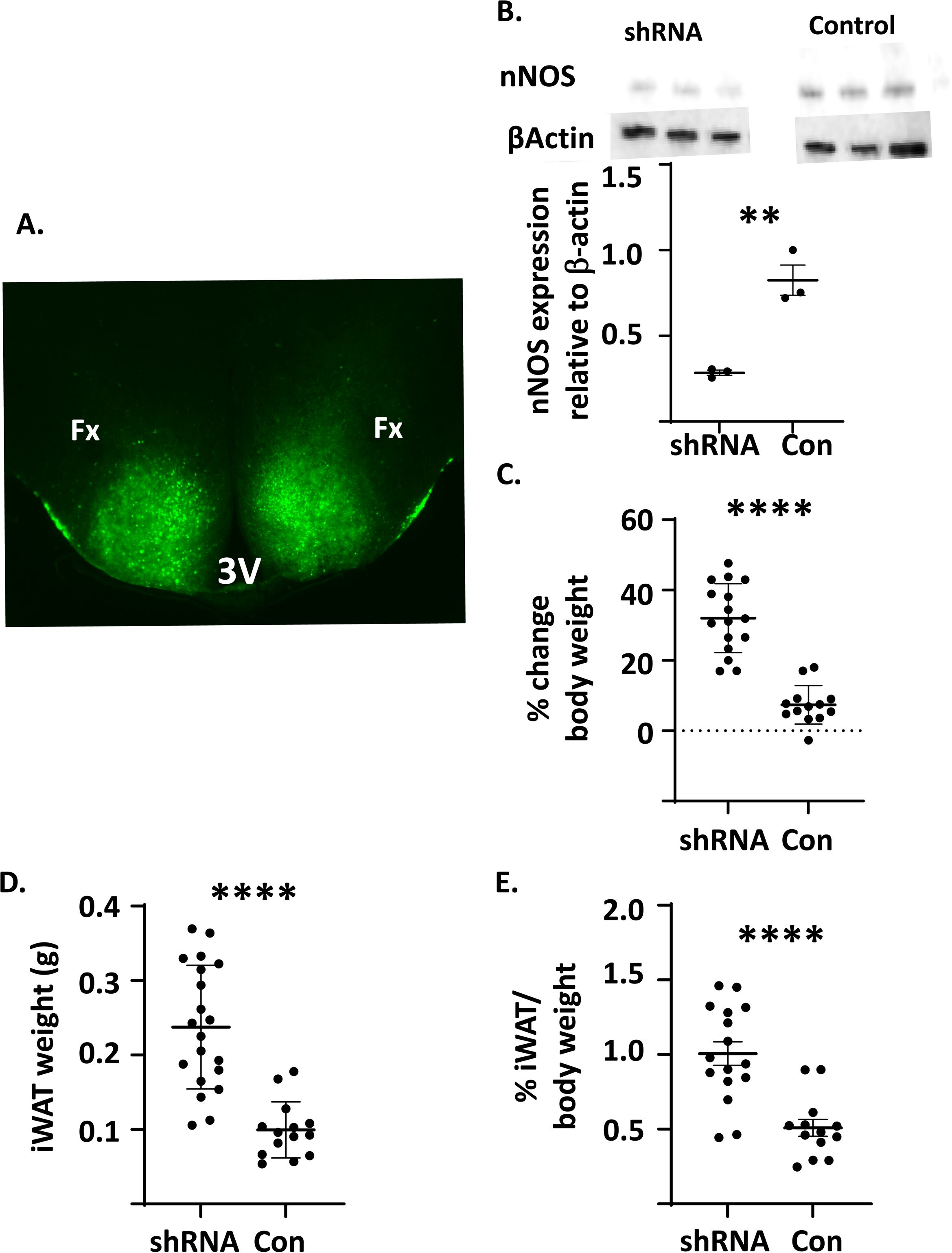
Reducing VMH nNOS expression increased body weight and adiposity. A. nNOS shRNA (green) injections were specific to the VMH. B. nNOS shRNA injection into the VMH reduced nNOS expression compared to that in mice injected with scrambled control AAV (con). C-E. 2 weeks post-injection VMH nNOS shRNA significantly increased the percent change in body weight (C), inguinal white fat weight (iWAT; D) and the percent of iWAT per gram body weight (E).

Mice injected with NOS1 shRNA into the VMH were fatter than scramble AAV injected controls. Body weights in shRNA injected mice increased by ∼30% at 2 weeks post-injection (Figure 3C). This was associated with an increase in inguinal fat weight and the percent of inguinal fat per body weight (Figure 3 D, E). BAT weight and vacuole size in inguinal fat were also elevated (Figure 4A). However, there were no differences in UCP1 expression in either BAT or inguinal fat in females or males (Figure 4 B, C). There was also no difference in intrascapular temperature in mice injected with shRNA compared to scrambled AAV injected controls (Figure 5A-C). Interestingly, shRNA injected mice showed a reduced activity index indicating a decrease in locomotion compared to controls (Figure 5 D-F). This was evident in both the dark and light cycle. There was no significant difference in activity between the light and dark cycle in either shRNA or scrambled AAV injected controls (Figure 5 G, H).

**Figure 4.**
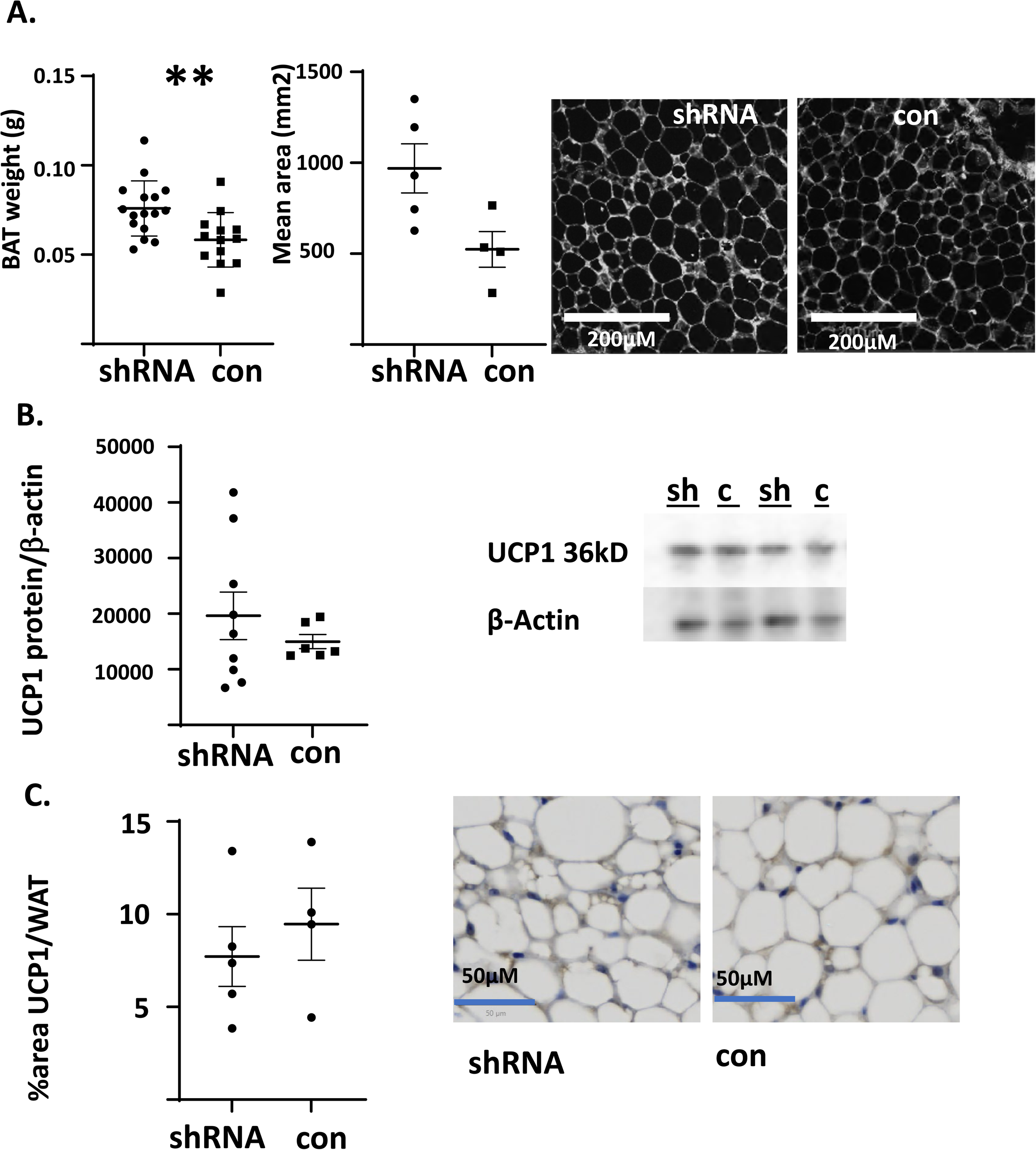
Reducing VMH nNOS expression did not alter BAT thermogenesis or WAT browning. A. Interscapular (i) BAT weight was increased by VMH nNOS shRNA relative to the control (con) AAV; however mean area of brown adipocytes was not changed. B. VMH nNOS shRNA (sh) did not alter iBAT uncoupling protein (UCP1) expression. C. VMH nNOS shRNA did not alter iWAT UCP1 expression. Left panel: UCP1 expression per area of iWAT was unchanged. Right panel: representative image of UCP1 (dark blue) relative to iWAT cells was unchanged.

**Figure 5.**
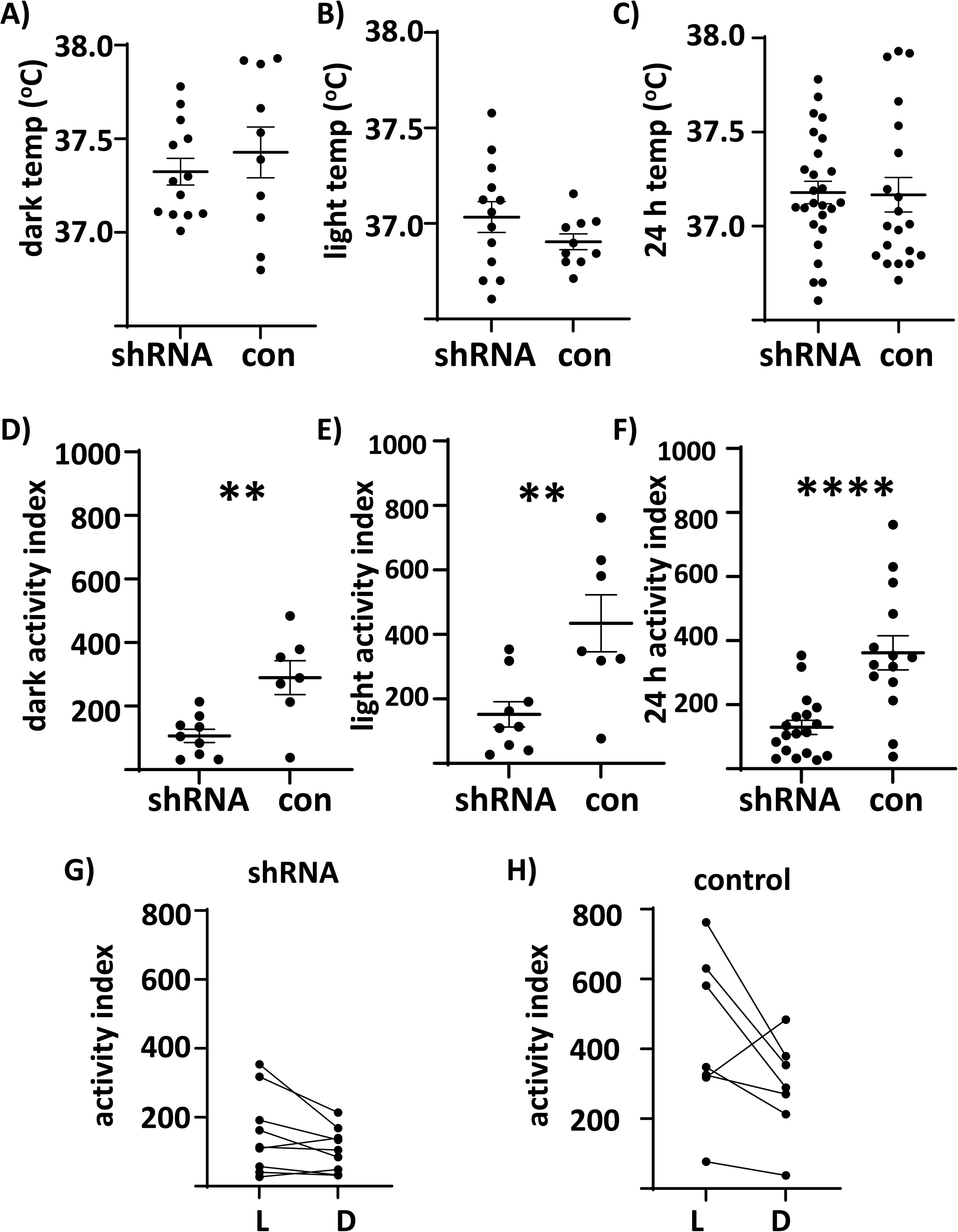
Reducing VMH nNOS expression reduced locomotor activity. A-C. VMH nNOS shRNA had no effect on interscapular temperature in the dark (A) or light (B) cycle, nor did it affect 24-hour temperature (C). D-E. VMH nNOS shRNA decreased the activity index in the dark (D) and light (E) cycle as well as decreasing the 24-hour activity index (F). G-H. There was no significant difference between light and dark cycle activity for either the shRNA or scrambled control AAV injected mice.

Blood glucose levels were significantly elevated in shRNA injected mice (Figure 6A). However, serum FFAs and epinephrine levels did not differ from scramble AAV injected controls (Figure 6 B, C). There was no correlation between glucose, FFAs and epinephrine and the % change in body weight.

**Figure 6.**
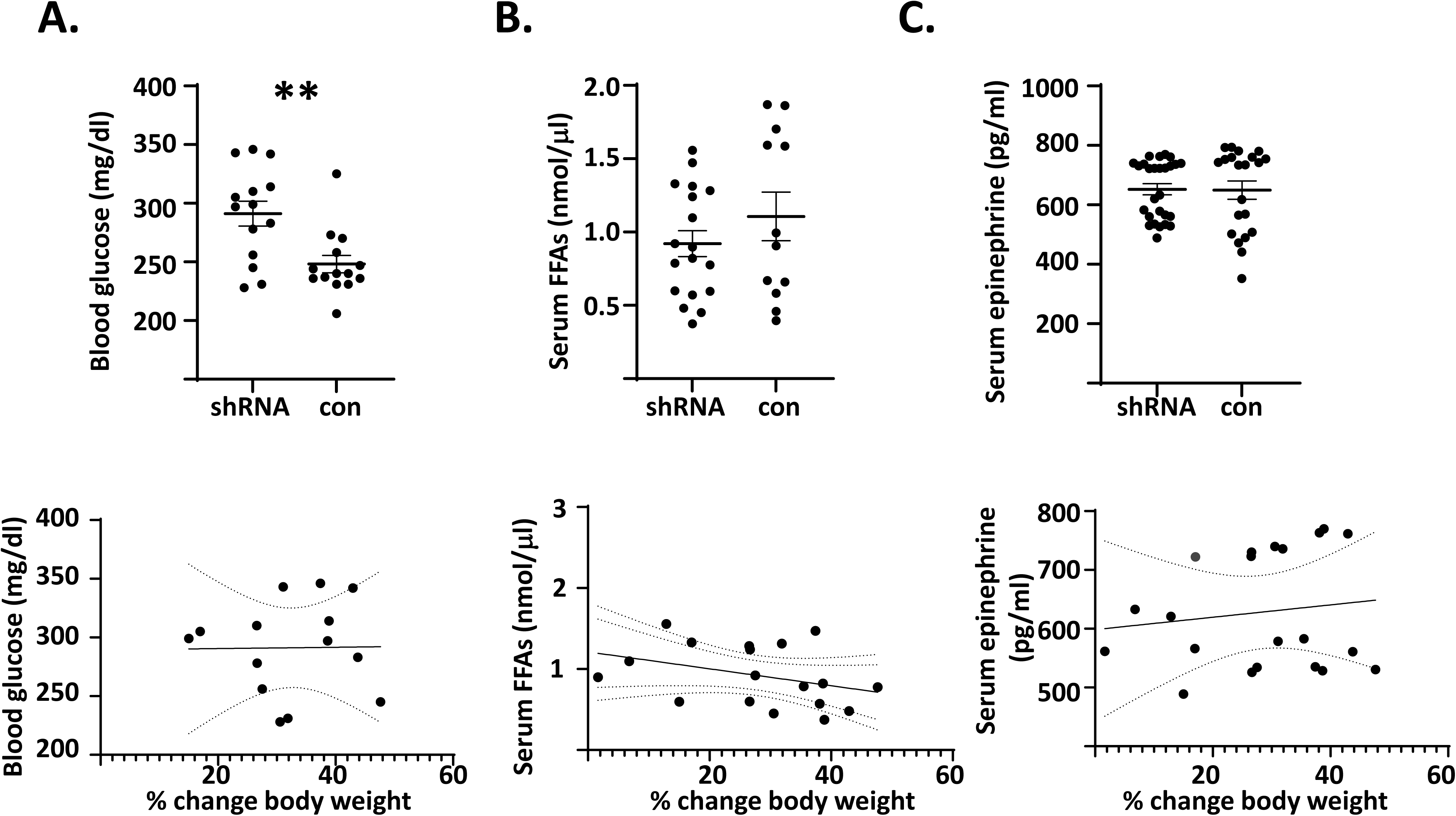
Reducing VMH nNOs expression reduced blood glucose levels. A. Blood glucose was significantly reduced in VMH shRNA injected mice. B-C. Serum free fatty acids and epinephrine were unaffected by VMH nNOS shRNA injection. There was no correlation between the percent change in body weight and blood glucose or serum free fatty acids and epinephrine levels.

### VMH neurons modulate glutamate and GABAergic currents onto orexin and non-orexin LH neurons, respectively

Given the lack of an nNOS-cre mouse we were unable to determine whether nNOS expressing VMH neurons modulated the activity of LH neurons. Instead, we injected a fluorescently labeled anterograde ChR2 into the VMH of female and male orexin-GFP mice and recorded from orexin and non-orexin LH neurons (Figure 7). The majority of LH orexin (Figure 7A) or non-orexin (Figure 7D) neurons showed no response to blue light stimulation. However, of the orexin neurons 9 out of 39 (∼23%) showed an inward current that was blocked by the glutamate receptor blocker, kynurenic acid (Figure 7B), while only 3 (∼7%) showed inward currents blocked by the GABA receptor blockers, bicuculline and saclofen (Figure 7C). In contrast, 5 of 21 (∼24%) LH non-orexin neurons (Figure 7E) had a response that was blocked by bicuculline (100uM) and saclofen indicating that they were GABAergic inputs (100uM) and only 1 of 21 (∼5%) had a response that was blocked by kynurenic acid (KA, 3mM; Figure 7F). Overall, synaptic currents due to VMH stimulation showed a higher amplitude and decay time in non-orexin vs orexin neurons (Fig 7 G, H).

**Figure 7.**
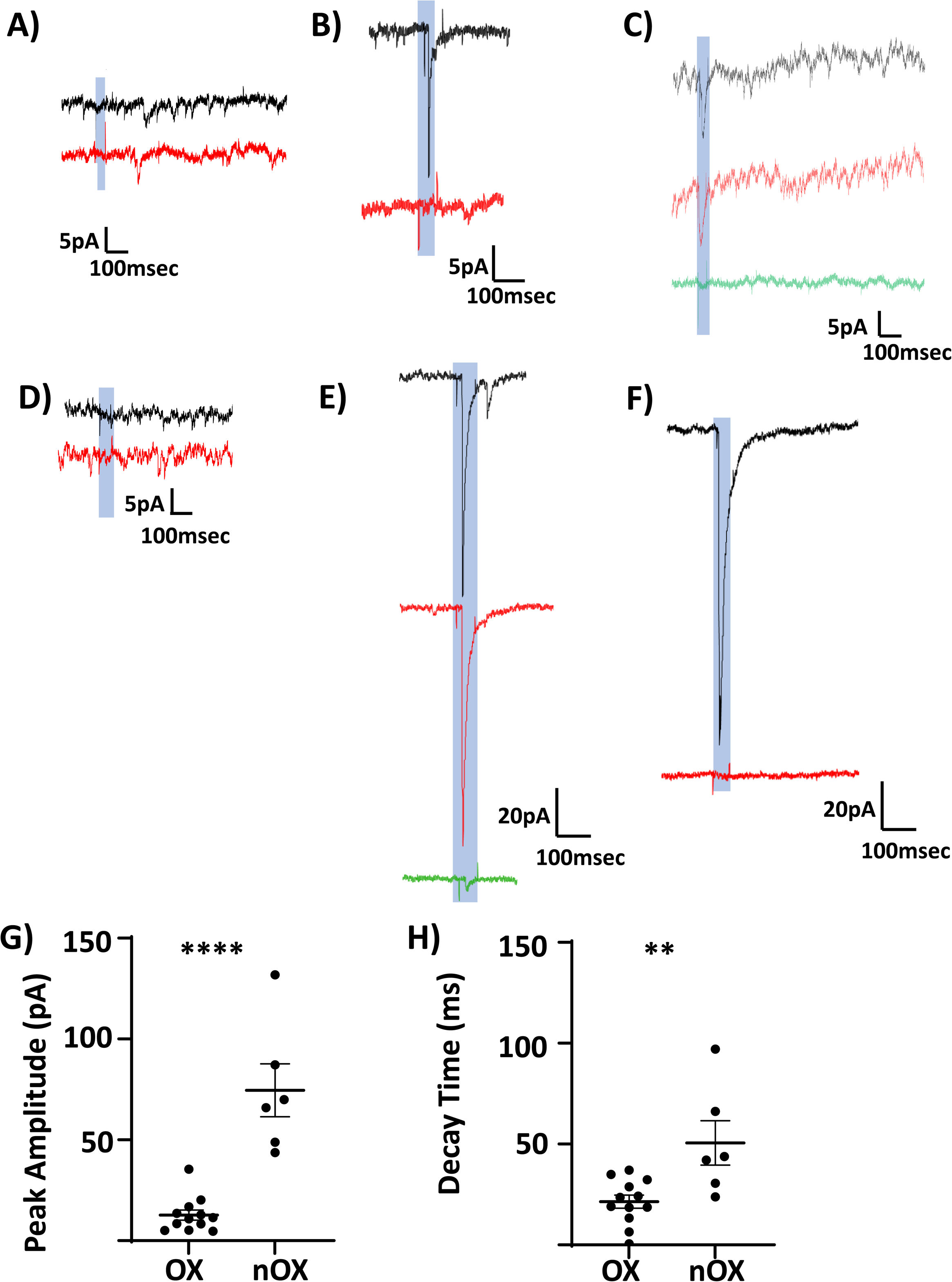
Voltage clamp recordings of LH neurons in brain slices from mice expressing green fluorescent protein (GFP) in orexin neurons. Mice had channelrhodopsin injected into the VMH and blue light was used to optogenetically activate neurons being recorded. A and D: most orexin and non-orexin neurons showed no response to blue light as indicated by representative traces. B. Representative trace of an orexin neuron exhibiting an inward current in response to blue light; this current (upper trace, black) was blocked by the glutamate receptor antagonist, kynurenic acid (lower trace, red). C. Representative trace of an orexin neuron exhibiting an inward current in response to blue light (upper trace, black); this current was blocked by the GABA inhibitors, bicuculline and saclofen (lower trace, green), but not by kynurenic acid (middle trace, red). E. Representative trace of a non-orexin neuron exhibiting an inward current in response to blue light (upper trace, black) that was not blocked by kyurenic acid (middle trace, red) but by the GABA inhibitors bicuculline and saclofen (lower trace, green). F. Representative trace of a non-orexin neuron exhibiting an inward current in response to blue light (upper trace, black) that was blocked by kynurenic acid (lower trace, red). G-H. Peak amplitude and decay time of the inward currents were greater in non-orexin vs orexin neurons.

## DISCUSSION

Our current and previous data show that vlVMH GI neurons require AMPK and nNOS for activation in low glucose^3,5,6,11^. Moreover, we now show that vlVMH-GI and -nNOS neurons project to the LH. However, reducing VMH nNOS expression had no effect on BAT thermogenesis or WAT browning. Instead, it resulted in a dramatic increase in body weight over 2 weeks that was associated with increased blood glucose and decreased locomotion. Additionally, of the orexin neurons that responded to VMH stimulation the majority showed glutamate currents indicating that the VMH stimulates orexin neurons. Together these data are not consistent with our original hypothesis that vlVMH nNOS-GI neurons mediate the orexin-dependent effects of estrogen on BAT thermogenesis and WAT browning. This is because the stimulatory effect of estrogen on orexin neurons, BAT and WAT depends on VMH AMPK inhibition^1^. We have previously shown that estrogen blunts activation of vlVMH GI neurons in low glucose by inhibiting AMPK^3^. Since these neurons require nNOS for activation^5,11^, we originally hypothesized that vlVMH nNOS-GI neurons inhibited orexin neurons and reducing VMH nNOS expression would thus increase BAT thermogenesis and WAT browning. However, this did not occur. Our results and the literature suggest that there are 2 populations of VMH AMPK dependent neurons; one of which inhibits the orexin neurons that facilitate BAT and WAT thermogenesis and another that stimulates the orexin neurons leading to increased physical activity and decreased glucose and body weight. Our data suggest the hypothesis that the vlVMH nNOS-GI neurons fall into the latter category of AMPK dependent VMH neurons.

Orexin neurons are related to a wide range of physiological actions including food and drug motivated behavior, arousal, sympathetic activation, and spontaneous physical activity^12,13^. Kotz and colleagues demonstrated that orexin neurons increase spontaneous physical activity and decrease blood glucose^4^. Our findings that 1) VMH nNOS shRNA increased body weight and blood glucose while decreasing locomotion; 2) most orexin cells responding to VMH stimulation received excitatory glutamatergic input and 3) vlVMH GI and nNOS neurons project to the LH are consistent with a role of vlVMH nNOS-GI neurons in stimulating physical activity and lowering glucose and body weight via excitation of orexin neurons. However, there are several caveats. First, we were unable to definitively determine that vlVMH nNOS-GI neurons excite orexin neurons. This is because we found cre recombinase expression in nNOS-cre mice was not restricted to nNOS expressing neurons. We used several methods to attempt to confirm expression (Figure 2). We evaluated nNOS and cre co-expression in 2 different colonies we established from separate founders obtained from Jackson laboratories (at least 1 year apart). These mice showed widespread cre-immunoreactivity that was not co-expressed with nNOS. Second, we bred both male and female mice obtained directly from Jackson laboratories to female or male mice, respectively, expressing cre-dependent GFP. These mice not only showed ectopic cre-expression in the brain but were visibly green. It is important to note that we used male nNOS-cre mice due to reports of issues in the female germline. Earlier papers from the laboratory that donated these mice convincingly demonstrated cre recombinase expression only in nNOS neurons^14^. The original strain was on the 129 background. Thus, the strain may have drifted when backcrossed onto the 6J background or during subsequent breeding. Non-specific cre recombinase expression has been reported previously in this strain^15^. Thus, for this study we used a non-specific anterograde ChR2 to evaluate the effects of VMH stimulation onto orexin neurons, which predominantly induced glutamate currents. Interestingly, VMH stimulation evoked predominantly GABA currents onto non-orexin neurons. LH orexin interneurons receive local GABA input so if the VMH were inhibiting these interneurons it would disinhibit the orexin neurons leading to increased excitation.

The second caveat is that we cannot be certain that nNOS shRNA specifically targets nNOS-GI neurons. However, we are reasonably confident that this is the case because of the almost complete overlap we have found between VMH NO producing neurons and GI neurons. That is, nNOS inhibitors blunt the activation of >95% of VMH GI neurons in low glucose and we did not observe any VMH GI neurons in mice lacking nNOS expression^5,11^. Moreover, using membrane potential and NO sensitive dyes we found that over 75% of VMH neurons that produce NO were GI neurons^6^. It should be noted that membrane potential dye underestimates the number of GI neurons because this method only detects responses above a fairly robust threshold. Importantly, here we confirm that vlVMH GI neurons specifically are nNOS dependent. Thus, there is a high likelihood that our results reflect a decreased activation of VMH nNOS-GI neurons, specifically. A final caveat is that it is impossible to constrain viral injections to just the ventrolateral portion of the VMH and so our in vivo results reflect nNOS-GI neurons in the central and likely the dorsomedial VMH as well.

In summary, we find that reduced expression of VMH nNOS dramatically and rapidly increases body weight and adiposity in female mice. This increased adiposity is associated with increased blood glucose levels and decreased locomotor activity. These effects may be due to stimulation of LH orexin neurons since we found that VMH nNOS- and -GI neurons project to the LH and that VMH stimulation enhances glutamatergic input onto LH orexin neurons. This would be consistent with observations that LH orexin neurons lower blood glucose and body weight by increasing physical activity^4^. It remains to be seen whether VMH shRNA has similar effects in males.

## ACKNOWLEDGEMENTS

Supported in part by R01DK103676 (VHR), R01DK069861 (VHR), and 1F31DK126433 (PMH). We are grateful to UID, Lake Villa, IL for allowing us to borrow their Mouse Matrix Home Cage Monitoring System.

## Notes

### Competing Interest Statement

The authors have declared no competing interest.

### Summary of Updates

updated author orchid and funding acknowledgement

## REFERENCES

1. . Martínez de Morentin PB, González-García I, Martins L, et al. Estradiol regulates brown adipose tissue thermogenesis via hypothalamic AMPK. Cell metabolism. 2014;20(1):41–53.

2. . Martins L, Seoane-Collazo P, Contreras C, et al. A Functional Link between AMPK and Orexin Mediates the Effect of BMP8B on Energy Balance. Cell Reports. 2016;16(8):2231–2242.

3. . Santiago AM, Clegg DJ, Routh VH. Estrogens modulate ventrolateral ventromedial hypothalamic glucose-inhibited neurons. Molecular Metabolism. 10// 2016;5(10):823–833.

4. . Zink AN, Bunney PE, Holm AA, Billington CJ, Kotz CM. Neuromodulation of orexin neurons reduces diet-induced adiposity. Int J Obes (Lond). Apr 2018;42(4):737–745.

5. . Murphy BA, Fakira KA, Song Z, Beuve A, Routh VH. AMP-activated Protein Kinase (AMPK) and Nitric Oxide (NO) regulate the glucose sensitivity of ventromedial hypothalamic (VMH) glucose-inhibited (GI) neurons. AJP - Cell Physiology. 2009;297(3):C750–C758.

6. . Canabal DD, Song Z, Potian JG, Beuve A, McArdle JJ, Routh VH. Glucose, insulin and leptin signaling pathways modulate nitric oxide (NO) synthesis in glucose-inhibited (GI) neurons in the ventromedial hypothalamus (VMH). *AJP - Regulatory*, Integrative and Comparative Physiology. 2007;292(4):R1418–R1428.

7. López M, Tena-Sempere M. Estradiol effects on hypothalamic AMPK and BAT thermogenesis: A gateway for obesity treatment? Pharmacology & Therapeutics.

8. . Sheng Z, Santiago AM, Thomas MP, Routh VH. Metabolic regulation of lateral hypothalamic glucose-inhibited orexin neurons may influence midbrain reward neurocircuitry. Molecular and Cellular Neuroscience. 9// 2014;62(0):30–41.

9. . Teegala SB, Sarkar P, Siegel DM, et al. Lateral hypothalamus hypocretin/orexin glucose-inhibited neurons promote food seeking after calorie restriction. Molecular Metabolism. 2023/08/02/ 2023:101788.

10. . Ting JT, Lee BR, Chong P, et al. Preparation of Acute Brain Slices Using an Optimized N-Methyl-D-glucamine Protective Recovery Method. Journal of visualized experiments : JoVE. Feb 26 2018(132):53825.

11. . Fioramonti X, Marsollier N, Song Z, et al. Ventromedial Hypothalamic Nitric Oxide Production Is Necessary for Hypoglycemia Detection and Counterregulation. Diabetes. 2010;59(2):519–528.

12. . Aston-Jones G, Smith RJ, Sartor GC, et al. Lateral hypothalamic orexin/hypocretin neurons: A role in reward-seeking and addiction. Brain Research. 2/16/ 2010;1314:74–90.

13. . Harris GC, Aston-Jones G. Arousal and reward: a dichotomy in orexin function. Trends in Neurosciences. 10// 2006;29(10):571–577.

14. . Leshan RL, Greenwald-Yarnell M, Patterson CM, Gonzalez IE, Myers MG. Leptin action through hypothalamic nitric oxide synthase-1-expressing neurons controls energy balance. Nat Med. 2012;18(5):820–823.

15. . Smith ACW, Scofield MD, Heinsbroek JA, et al. Accumbens nNOS Interneurons Regulate Cocaine Relapse. The Journal of Neuroscience. 2017;37(4):742–756.

